# Modeling the relationship between estimated foliar fungicide use and soybean yield losses due to foliar fungal diseases in the United States

**DOI:** 10.1101/744581

**Authors:** Ananda Y. Bandara, Dilooshi K. Weerasooriya, Shawn P. Conley, Carl A. Bradley, Tom W. Allen, Paul D. Esker

## Abstract

Fungicide use in the United States to manage soybean diseases has increased in recent years. The ability of fungicides to reduce disease-associated yield losses varies greatly depending on multiple factors. Nonetheless, historical data are useful to understand the broad sense and long-term trends related to fungicide use practices. In the current study, the relationship between estimated soybean yield losses due to selected foliar diseases and foliar fungicide use was investigated using annual data from 28 soybean growing states over the period of 2005 to 2015. At a national scale, a significant quadratic relationship was observed between total estimated yield losses and total fungicide use (R^2^ = 0.123, *P* < 0.0001) where yield losses initially increased, reached a plateau, and subsequently decreased with increasing fungicide use. The positive phase of the quadratic curve could be associated with insufficient amount of fungicides being used to manage targeted diseases, application of more-than-recommended prophylactic fungicides under no/low disease pressure, application of curative fungicides after economic injury level, and reduced fungicide efficacy due to a variety of factors such as unfavorable environmental conditions and resistance of targeted pathogen populations to the specific active ingredient applied. Interestingly, a significant quadratic relationship was also observed between total soybean production and total foliar fungicide use (R^2^= 0.36, *P* < 0.0001). The positive phase of the quadratic curve may suggest that factors like plant physiological changes, including increased chlorophyll content, photosynthetic rates, water use efficiency, and delayed senescence that have been widely reported to occur after application of certain foliar fungicides could have potentially contributed to enhanced yield. Therefore, the current study provides evidence of the potential usefulness of foliar fungicide applications to mitigate soybean yield losses associated with foliar diseases and their potential to positively impact soybean production/yield at national and regional scales although discrepancies to the general trends observed at national and regional scales do prevail at the local (state) level.

## INTRODUCTION

Soybean [*Glycine max* (L.) Merrill] is a key agricultural commodity in the United States and has been cultivated on 34.7 million hectares on average annually between 2015 and 2019 (USDA-NASS). Similar to the production of other economically important crops, numerous abiotic and biotic stressors like adverse weather, variation in soil characteristics, diseases, insects, and weeds present enormous challenges to soybean production [1, 2]. Soybean diseases are detrimental to production due to their deleterious effects on yield. In the U.S., the average annual disease-associated soybean yield losses are approximately 11% [3]. However, the relative importance of diseases and concomitant yield losses vary both temporally and spatially. For example, total yield losses due to diseases in 2012 was estimated to be 10.07 million metric tons while in 2014 it was 13.94 million metric tons [4]. Among various soybean foliar diseases, Septoria brown spot, caused by *Septoria glycines* Hemmi, and frogeye leaf spot, caused by *Cercospora sojina* Hara, are the most common [1, 5–8] and are also considered to be important yield limiting diseases in soybean [9]. The losses caused by Septoria brown spot range from 196 to 293 kg ha^−1^ [6]. Septoria brown spot can cause up to 2,000 kg ha^−1^ loss in high-yield soybean production systems (>5,000 kg ha^−1^) [10]. Frogeye leaf spot can result in yield losses from 10 to 60% [11] and seed weight reductions up to 29% [12].

Different management strategies are deployed either individually or in an integrated manner to reduce the losses caused by foliar fungal diseases in soybean production systems. Among these, the use of foliar-applied fungicides has been an important tactic. Fungicide use in soybean has risen dramatically since 2005 [13]. Several reasons were given to explain this increase including: increased availability of fungicides for use on soybean, improved awareness of soybean diseases, the initial observation of soybean rust in North America and the resultant production of specific chemistries to manage this disease that were not widely used, increased soybean commodity price, and promotion of certain fungicides by the manufacturers for their potential physiological benefits that may increase soybean yield even in the absence of disease, a phenomenon in which the term “plant health” has been coined [14, 15].

The quinone-outside inhibitor (QoI; strobilurin) class of fungicides (Fungicide Resistance Action Committee [FRAC] group 11) are commonly used to manage foliar diseases of soybean and these act by binding with complex III of the mitochondrial respiration pathway [16]. Additionally, the demethylation inhibitor (DMI; triazole) class of fungicides (FRAC group 3) are also used in soybean and this class of fungicides inhibit ergosterol biosynthesis by fungi [17]. Recently, active ingredients from the succinate dehydrogenase inhibiting (SDHI; FRAC group 7) class of fungicides were introduced for management of foliar soybean diseases. Similar to QoI fungicides, SDHI fungicides are classified as respiration inhibitors. However, instead of complex III, SDHI fungicides bind at complex II in the mitochondrial respiration pathway [17]. In general, these fungicide groups possess broad-spectrum activity on foliar fungal soybean diseases including Septoria brown spot and frogeye leaf spot [18]. The fungicides within these specific chemical classes can generally be purchased as stand-alone fungicides, especially those products designated as either DMI or QoI. However, stand-alone fungicide products consisting of SDHIs are currently not available and are included as a pre-mix fungicide that contains either one of the other classes (either DMI or QoI) or both of the classes as a three-way fungicide product. The current fungicide production trend from chemical manufacturers is to provide products that contain multiple modes of action to help reduce the development of fungicide resistance. In general, and to more broadly classify the chemical classes as outlined above, following the initial observation of soybean rust in the contiguous U.S., fungicide products were broadly categorized as either curative (DMI) and preventive (QoI and also SDHI).

Although foliar fungicides have extensively been used for soybean production, the extent to which yield losses can actually be mitigated with fungicide application and the subsequent economic return is often questioned. While fungicides are reported to reduce the yield losses when diseases are present [19, 20], the impact of fungicide application on yield in the absence of disease, i.e., the plant health scenario, are inconsistent. Several studies have demonstrated no significant increase in soybean yield with fungicide applications in the absence of disease [20–23], while other studies suggested that yield increases can occur with foliar fungicide application even in the absence of disease [7, 23–25]. Therefore, the economic return following a fungicide application does not intuitively follow a linear trend due to its apparent dependency on multiple factors such as disease pressure, class of fungicide being used (i.e., active ingredient), time of application (growth stage of the plant), and environmental conditions [19, 26, 27].

Widespread fungicide use can ultimately lead to an increased risk of selecting fungicide-resistant strains out of the targeted pathogen population. Fungicide resistance is an issue increasing in importance across soybean production areas in the U.S. as a result of automatic fungicide applications at specific growth stages, as well as fungicide applications with specific fungicide classes where the goal is a curative response [28–31]. Currently, QoI fungicide resistance has been reported for several soybean pathogens in the U.S., including *C. sojina*, in Illinois, Tennessee [32], South Dakota [33], and Mississippi [29]. Zhang et al [31] recently reported QoI resistant *C. sojina* isolates from 14 states including Alabama, Arkansas, Delaware, Illinois, Indiana, Iowa, Kentucky, Louisiana, Mississippi, Missouri, North Carolina, Ohio, Tennessee, and Virginia. Additionally, the fungi responsible for causing Cercospora leaf blight (*C*. cf. *flagellaris*, *C. kikuchii* (Tak. Matsumoto & Tomoy.) M.W. Gardner and *C*. cf. *sigesbeckiae*) have been reported to exhibit resistance to QoI fungicides throughout Louisiana [28]. Moreover, additional anecdotal, unpublished reports of resistance within populations of *S. glyinces* and *Corynespora cassiicola* (Berk. & M.A. Curtis) C.T. Wei, the causal organism of target spot of soybean have recently been made.

In the current paper, we investigate long-term fungicide use patterns and the relationship with soybean yield and the resulting foliar diseases that cause losses. Our primary spatial grain was at the state level, although regional and national level trends were also explored. While numerous individual experiments have been conducted to address the aforementioned issues, a more comprehensive analysis with long term historical data (estimated fungicide use and soybean yield losses as a result of diseases) is currently lacking. Thus, our objectives for this study were to (i) investigate the relationship between foliar fungicide use in the U.S. and estimated yield losses due to foliar diseases, and (ii) investigate the relationship between foliar fungicide use in the U.S. and soybean production/yield at national, regional, and state levels. Findings of this study will aid in informed decision making on spatiotemporally sensitive, economically viable, and environmentally sound use of fungicides to manage soybean fungal diseases in the U.S. Furthermore, results will also provide useful insights into how research, policy, and educational efforts should be prioritized in soybean disease management using fungicides.

## MATERIALS AND METHODS

### Fungicide use data

Annual state-level foliar fungicide use estimates (in Kg of active ingredient) for soybean were obtained from the Pesticide National Synthesis Project webpage (https://water.usgs.gov/nawqa/pnsp/usage/maps/county-level/StateLevel/HighEstimate_AgPestUsebyCropGroup92to16.txt). Foliar fungicides applied to soybean during the period between 2005 and 2015 were considered for this study. The time period was based upon the availability of fungicide use data spanning 28 soybean growing states (AL, AR, DE, FL, GA, IA, IL, IN, KS, KY, LA, MD, MI, MN, MO, MS, NC, ND, NE, OH, OK, PA, SC, SD, TN, TX, VA, WI). Fungicide use data were also classified based on each region where northern states considered for this study included IL, IN, IA, KS, MI, MN, NE, ND, OH, PA, SD, and WI while southern states included AL, AR, DE, FL, GA, KY, LA, MD, MS, MO, NC, OK, SC, TN, TX, and VA. The classification of states into regions was based on the two groups of soybean pathologists collecting disease loss estimate data, NCERA-137 (North Central Extension and Research Activity for Soybean Diseases) and the Southern Soybean Disease Workers.

To compute the fungicide use per unit area within each state (in grams per hectare), the amount provided in the database (in kg) was first converted to grams (g). The soybean planting and harvesting area was retrieved from USDA-NASS database (https://quickstats.nass.usda.gov) for individual states from 2005 to 2015. Fungicide use values (in g) were divided by respective state-wide total soybean (i) planted number of hectares and (ii) harvested number of hectares separately to decide the most appropriate type of explanatory variable (g of fungicide per unit hectarage planted versus g of fungicide per unit hectarage harvested) for use in the study. A simple linear regression analysis showed that two variables were linearly and positively related to each other (R^2^ = 0.9987, *P* < 0.0001, *y = 0.965x + 0.211*), indicating a high similarity between the two variables. As such, for this study, we report the fungicide concentration in grams of fungicide per harvested hectare (here after mentioned as g/ha).

### Yield loss data

Historical soybean yield loss estimates were gathered from soybean Extension specialists and researchers. We considered the soybean losses for the same periods where foliar fungicide data were also available. Soybean losses spanned the same 28 soybean growing states as indicated above. The methodology used to collect and report soybean disease losses have been previously described [4]. Briefly, a spreadsheet was circulated annually to plant pathologists with soybean responsibilities and they provided estimates of the losses associated with a defined set of diseases (n=23). However, for the purposes of this study we focused on the results related to foliar diseases caused by fungi that could be effectively managed by foliar fungicide application. The methods employed within each state differed with regards to the specific method for estimating losses; however, in general, some of the methods employed were based on each individual’s evaluation of cultivar trials, fungicide efficacy plots, specific troubleshooting or field calls, queries of Extension personnel within counties/parishes, statewide plant disease surveys, or plant disease diagnostic laboratory databases.

Given that the historical yield loss data were provided in the form of losses in metric tons (MT) of production, to calculate the loss per soybean disease, we first calculated the loss as a percentage based on overall production (in MT) per state and year using USDA-NASS data. We then calculated the overall loss (as a percentage) due to soybean diseases using Padwick’s calculation [40], which is:

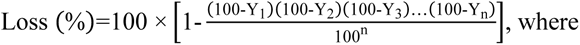

Y_1_, Y_2_, Y_3_, Y_n_, represent the percentage loss due to disease 1, 2, 3, through n, respectively. To estimate the loss due to diseases in terms of yield, we used the average soybean yield per state and year, from which we estimated the yield in the absence of diseases (the percentage loss estimated using Padwick’s calculation). The difference between the state average yield and the estimated yield in the absence of diseases was considered as the loss.

### Derivation of soybean yield, harvest, and production zones

The following categorical variables to be used in the factor analysis with mixed data (FAMD) and analysis of variance (ANOVA) (see below for details) were created: (i) Yield zone (1 to 4), based on USDA-NASS estimates at the state level comparing yield (MT/HA) with all state by year combinations, (ii) Harvest zone (1 to 4), based upon USDA-NASS estimates at the state level comparing harvested area (HA) with all state by year combinations, and (iii) Production zone (1 to 4), based upon USDA-NASS estimates at the state level comparing total production (MT) with all state by year combinations. Data points within the minimum to first quartile were classified as Zone 1. Similarly, data points from the first quartile to median, median to third quartile, and > third quartile were classified as zones 2, 3, and 4, respectively. Note that the zones were not solely defined based on geography, in this case state, and are a function of time (temporal scale). As such, the zone of a given data point was relative to the other data points (in terms of yield, harvest area, or total production) within the database. As yield, harvest area, and production within a given state fluctuated over time, the zone classification for a given state varied based on the year. The yield, harvest, and production zones corresponding to foliar fungicide data were therefore derived using soybean yield, harvest, and production data from 2005 to 2015.

### Fungicides and their targeted diseases considered

Based on data available in the fungicide and yield loss databases combined with soybean fungicide efficacy summarized by Extension plant pathologists on an annual basis, we concentrated on specific diseases for this study. Foliar fungicides (n=15) included the following active ingredients within several specific chemical classes as defined by the FRAC: QoIs (FRAC code 11) = azoxystrobin, fluoxastrobin, picoxystrobin, pyraclostrobin, trifloxystrobin; DMIs (FRAC code 3) = cyproconazole, difenoconazole, flutriafol, propiconazole, prothioconazole, tebuconazole, tetraconazole; chloronitrile (FRAC code M 05) = chlorothalonil; SDHI (FRAC code 7) = fluxapyroxad; and methyl benzimidazole carbamate (MBC) (FRAC code 1) = thiophanate-methyl. Although azoxystrobin, pyraclostrobin, and trifloxystrobin have uses as seed-applied fungicides, they were considered as foliar fungicides for this study as they are predominantly used to manage foliar diseases of soybean. The targeted diseases for the foliar fungicides listed above included anthracnose (caused by *Colletotrichum truncatum* (Schwein.) Andrus & W.D. Moore and several related species), Cercospora leaf blight (purple seed stain: *Cercospora flagellaris, C. kikuchii, C. sigesbeckiae*), frogeye leaf spot (*Cercospora sojina*), Rhizoctonia aerial blight (*Rhizoctonia solani* J.G. Kühn), Sclerotinia stem rot (White mold: *Sclerotinia sclerotiorum* (Lib.) de Bary), Septoria brown spot (*Septoria glycines*), and soybean rust (*Phakopsora pachyrhizi* Syd. & P. Syd.).

### Determination of the relationship between fungicide use and yield losses due to diseases

The PROC GLM procedure in SAS (version 9.4, SAS Institute, 2017, Cary, NC) was used to analyze the strength of relationships (as indicated by the coefficient of determination: R^2^) at the national, regional, state levels, as well as at temporal scale (on an annual basis). Linear and second order polynomial (=quadratic) curves were fitted to determine the most realistic relationship among the different variables. In cases where both linear and quadratic relations were significant, the quadratic curve was selected for the interpretation purpose. The higher order (third or more) polynomial curves were not fitted due to the lack of interpretability despite the possible significant model *P*-values associated with these curves. The analyses were conducted to examine total fungicide use (MT) and total yield loss (1,000 MT), as well as total fungicide use per unit harvest area (g/ha) and total yield loss per unit area (kg/ha). An initial analysis was conducted for the whole data set (all states, all years) to investigate the national trend over time. Subsequent analyses were conducted to evaluate different trends at the regional and state level, as well as in temporal scale. In addition, similar analyses (as indicated above) were performed to investigate the relationship between fungicide use and soybean production/yield.

### Factor Analysis with Mixed Data (FAMD)

FAMD is a principal component method to analyze a data set containing both quantitative and qualitative variables [35]. FAMD makes it possible to analyze the similarity between individuals (individual data points) by taking into account mixed-variable types. With this analysis, quantitative and qualitative variables are normalized in order to balance the impact of each set of variables. The packages FactoMineR version 1.41 (for the analysis) and factoextra (for data visualization) in R (version 3.5.1) were used for FAMD analysis. Here, total foliar fungicide use in grams of active ingredient (on a per hectare (ha) basis) was used as a quantitative variable while the year, state, region, soybean yield zone, harvest zone, and production zones, were incorporated as qualitative variables.

### Analysis of variance (ANOVA)

To investigate the main effects of yield, harvest, and production zones on total fungicide use (per ha basis), ANOVA was conducted using the PROC GLIMMIX procedure in SAS (version 9.4, SAS Institute, Cary, NC) at the 5% significance level. Restricted maximum likelihood (REML) was used to compute the variance components. Degrees of freedom for the denominator of F tests were computed using the Kenward-Roger option. Studentized residual plots and Q-Q plots were respectively used to assess the assumptions of identical and independent distribution of residuals, and their normality. Appropriate heterogeneous variance models were fitted whenever heteroskedasticity was observed by specifying a “*random residual/group = x*” statement (where *x* = factor under consideration, ex: yield zone). The Bayesian information criterion (model with the lowest BIC) was used to select the best fitting model (between homogenous variance vs heterogeneous variance). Mean separation was performed with adjustments for multiple comparisons using the Tukey-Kramer test.

## RESULTS

### Temporal fluctuation of soybean fungicide use in the United States

Considering total fungicide use (in both MT and g/ha) across 28 soybean growing states, the greatest foliar fungicide use was recorded in 2007 with the lowest recorded use in 2006 (Fig. 1A). A 63.5% decrease in foliar fungicide use on a per ha basis was evident from 2007 to 2008. The percentage use increment from 2006 to 2015 was 317% for total fungicide use in MT and 252% for total fungicide use in g/ha, respectively. Despite the annual variation, the total concentration of foliar fungicides used in 28 states showed a general increasing trend from 2005 to 2015.

**Fig 1.**
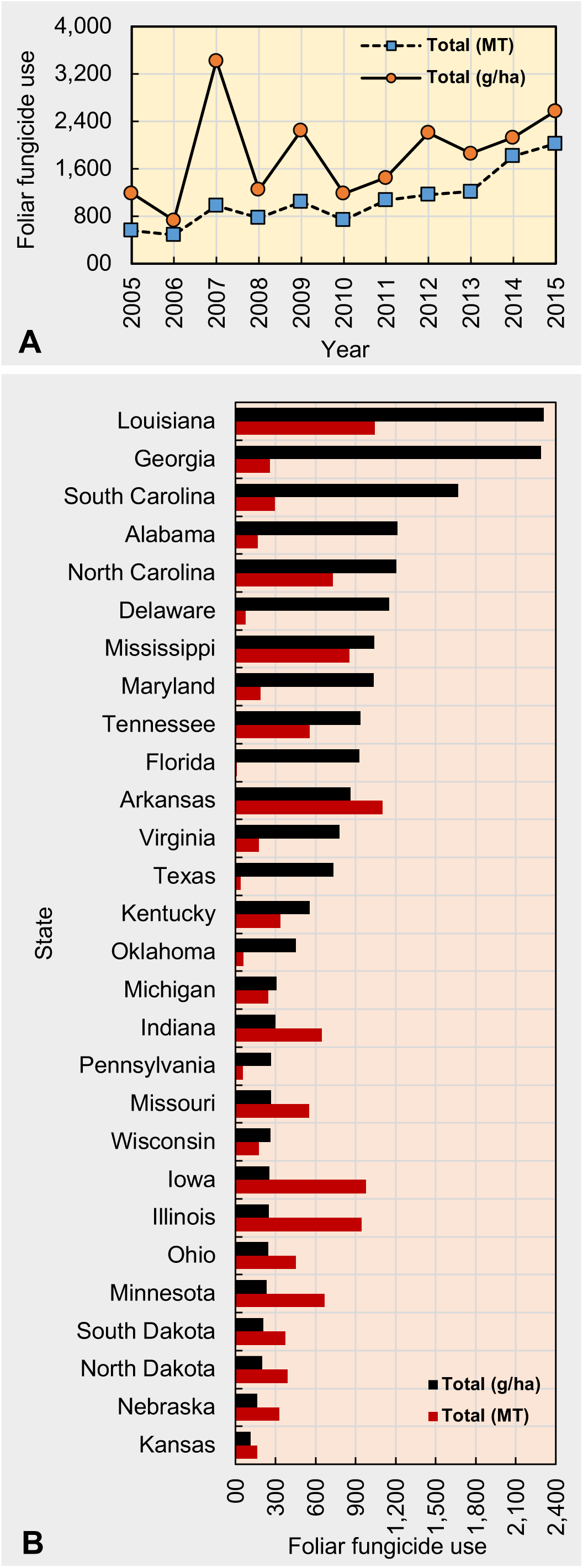
Spatiotemporal foliar fungicide use patterns in the United States. Temporal fluctuation for foliar fungicide use during 2005 to 2015 across all states considered (A) and state-wide use of cumulative foliar fungicides from 2005 to 2015 (B). Fungicides included: quinone outside inhibitors = azoxystrobin, fluoxastrobin, picoxystrobin, pyraclostrobin, trifloxystrobin; demethylation inhibitors = cyproconazole, difenoconazole, flutriafol, propiconazole, prothioconazole, tebuconazole, tetraconazole; methyl benzimidazole carbamates = thiophanate-methyl; multi-site mode of action = chlorothalonil; and succinate dehydrogenase inhibitors = fluxapyroxad.

### Spatial fluctuation of soybean fungicide use in the United States

Over an 11-year period, between 2005 and 2015 on a per hectare basis, Louisiana reported the greatest foliar fungicide use (2,309 g) while Kansas reported the lowest (114 g) (Fig. 1B). In terms of the total foliar fungicide use (in MT), Florida recorded the lowest (9.7 MT) while Arkansas reported the greatest (1,103.7 MT).

When considered regionally, the total use (MT) of foliar fungicides was 18.7% greater in the southern states (6,451.3 MT) compared to northern states (5,431.2 MT) (Fig. 2). Similarly, per hectare total use (g/ha) of foliar fungicides was 521% greater in the southern states (17,437.2 g/ha) compared to the northern states (2,805.7 g/ha) (Fig. 2).

**Fig 2.**
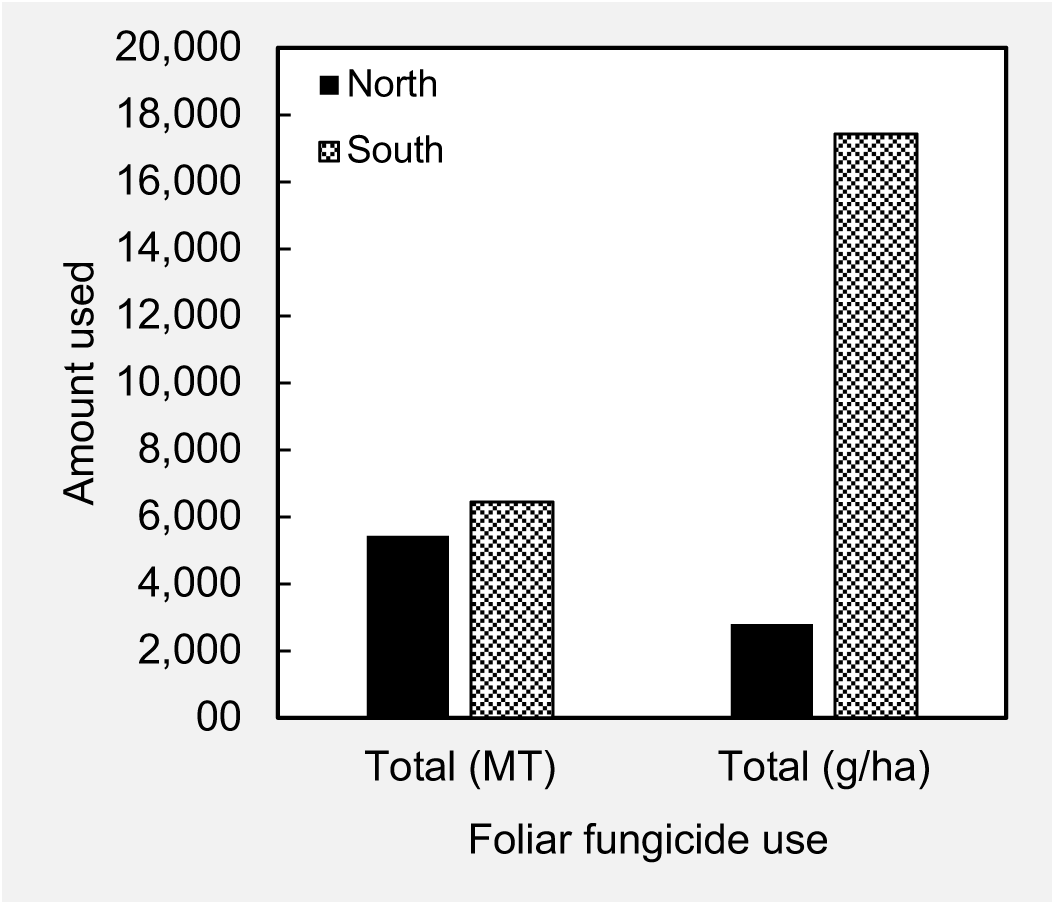
Total foliar fungicide use (from 2005 to 2015) by region. Northern states = IL, IN, IA, KS, MI, MN, NE, ND, OH, PA, SD, and WI; Southern states = AL, AR, DE, FL, GA, KY, LA, MD, MO, MS, NC, OK, SC, TN, TX, and VA. Fungicides included: quinone outside inhibitors = azoxystrobin, fluoxastrobin, picoxystrobin, pyraclostrobin, trifloxystrobin; demethylation inhibitors = cyproconazole, difenoconazole, flutriafol, propiconazole, prothioconazole, tebuconazole, tetraconazole; methyl benzimidazole carbamates = thiophanate-methyl; multi-site mode of action = chlorothalonil; and succinate dehydrogenase inhibitors = fluxapyroxad.

### Preventive vs curative fungicides

In general, the QoI class of fungicides, commonly referred to as strobilurins are used as preventative fungicides while DMI (or triazoles) are used as curative fungicides. Temporal fluctuations (summed across states) showed that the use of both types of fungicides increased from 2005 to 2015 (Fig. 3A). The amount of preventive and curative fungicides used in 2015 were 3.34 and 4.2-fold greater compared to their use in 2005. The use of QoI fungicides, representing = ∑ azoxystrobin, fluoxastrobin, picoxystrobin, pyraclostrobin, and trifloxystrobin, was greater compared to curative fungicides representing = ∑ cyproconazole, difenoconazole, propiconazole, prothioconazole, tebuconazole, and tetraconazole for any given year. Spatially, the greatest and lowest QoI fungicide use, summed across years, was recorded in Iowa and Florida, respectively, while the greatest and lowest DMI fungicide use was recorded in Illinois and Florida, respectively (Fig. 3B). In general, QoI fungicide use was greater compared to DMI fungicides except in a few states (Alabama, Delaware, Georgia, South Carolina, and South Dakota).

**Fig 3.**
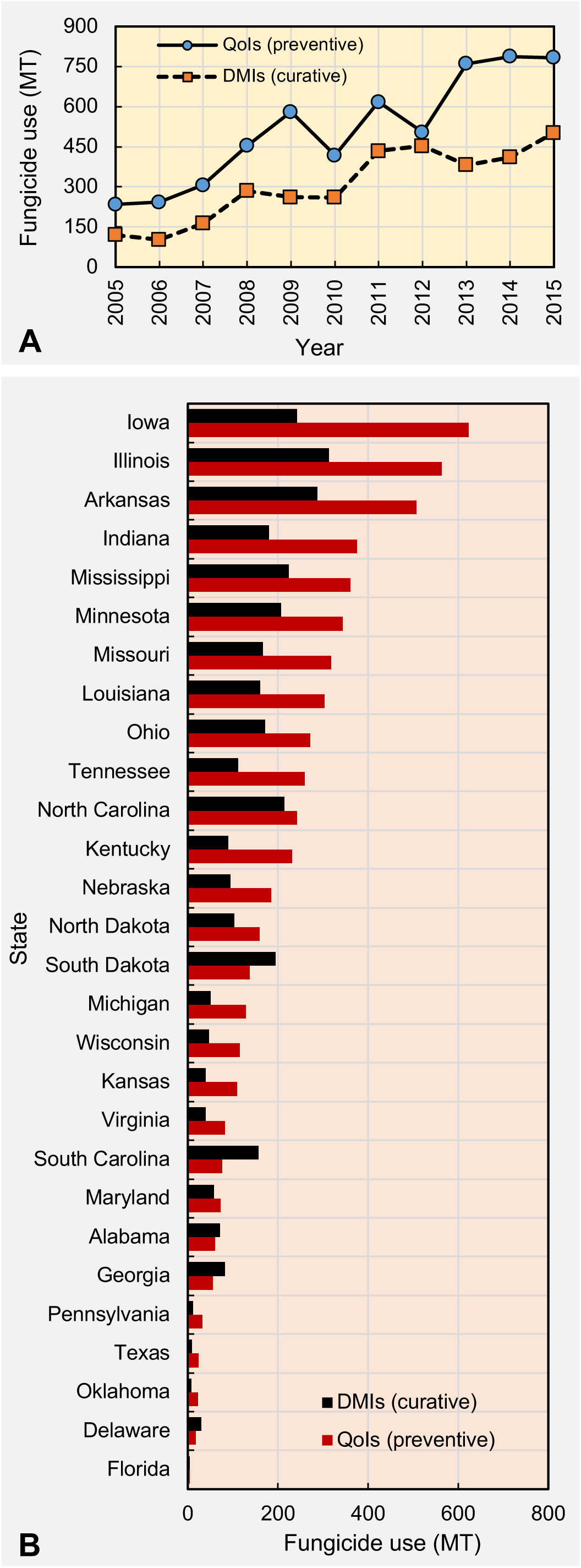
Temporal fluctuation (A) and state-wide variation (B) in the amount of preventive and curative foliar fungicide application use in the United States. Preventive fungicides = quinone outside inhibitors (QoIs) = ∑ azoxystrobin, fluoxastrobin, picoxystrobin, pyraclostrobin, and trifloxystrobin. Curative fungicides = demethylation inhibitors (DMIs) = ∑ cyproconazole, difenoconazole, flutriafol, propiconazole, prothioconazole, tebuconazole, and tetraconazole.

### Relationship between annual soybean yield losses and annual fungicide use at national and regional levels

Table 1 shows the coefficient of determination (R^2^) and corresponding *P*-values for the linear and quadratic relationships between yield losses and fungicide use at the national and regional levels. The analysis of the complete data set at the national level revealed a significant quadratic relationship between total yield losses due to foliar diseases (1,000 MT) and total foliar fungicide use (MT) during the period from 2005 to 2015 (Fig. 4A). Based on this relationship, the total yield losses initially increased, reached a plateau, and subsequently decreased with increasing fungicide use. When the losses (kg) and fungicide use (g) were considered on a per hectare basis, there was no significant linear or quadratic relationship (Fig. 4B).

**Fig 4.**
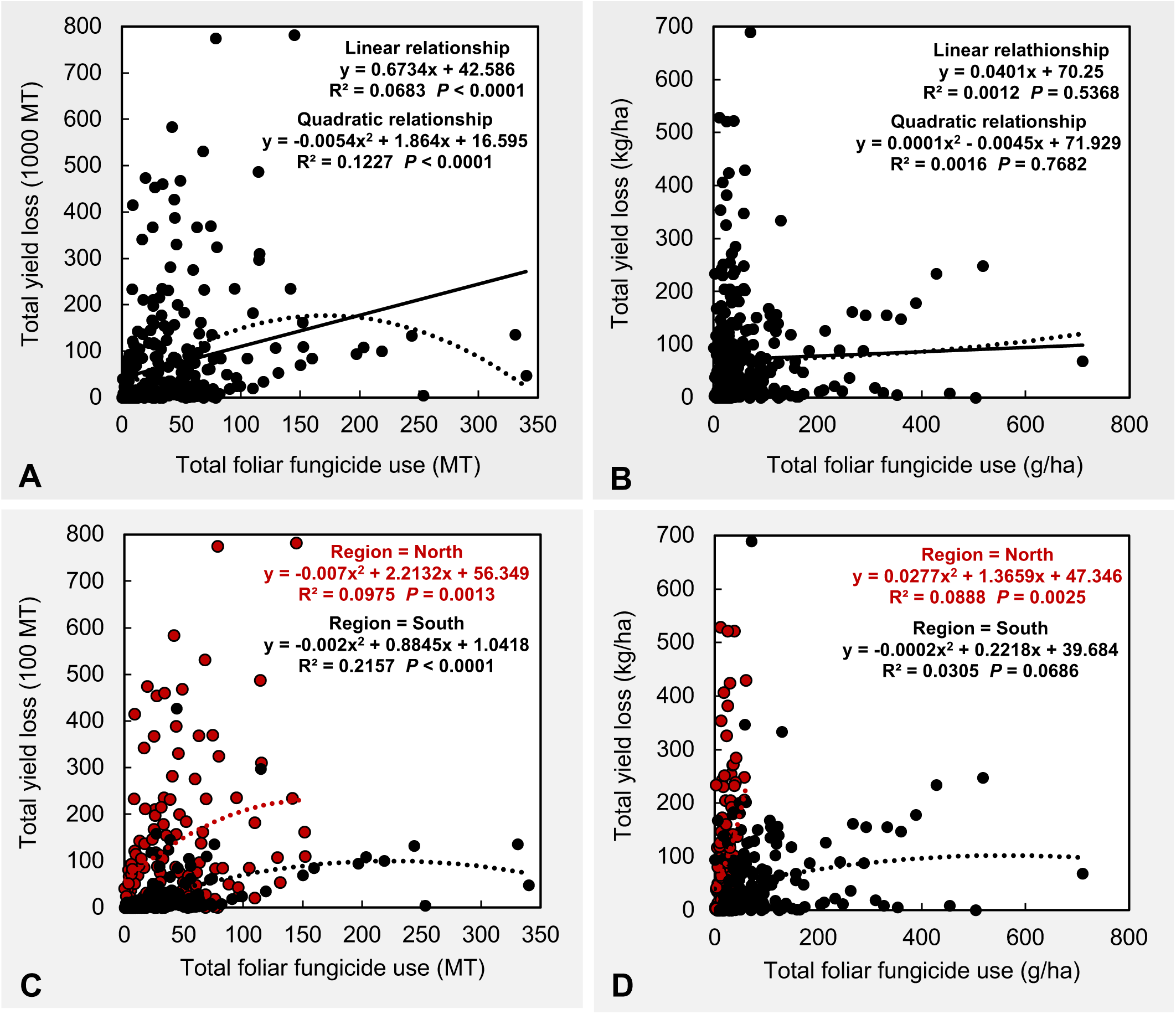
Scatter plots presenting the national scale linear/quadratic relationship between (A) total soybean yield losses due to foliar diseases (1,000 MT) and total foliar fungicide use (MT), (B) per hectare total soybean yield losses due to foliar diseases (in kg) and per hectare total foliar fungicide use (g), and regional scale quadratic relationship between (C) total soybean yield losses due to foliar diseases (1,000 MT) and foliar fungicide use (MT), and (D) per hectare total soybean yield losses due to foliar diseases (in kg) and per hectare total foliar fungicide use (g). Each data point represents a state in a given year. All figures contain data during 2005 to 2015 from 28 soybean growing states including AL, AR, DE, FL, GA, IA, IL, IN, KS, KY, LA, MD, MI, MN, MO, MS, NC, ND, NE, OH, OK, PA, SC, SD, TN, TX, VA, and WI. Foliar diseases include anthracnose, Cercospora leaf blight (purple seed stain), frogeye leaf spot, Rhizoctonia aerial blight, Sclerotinia stem rot (White mold), Septoria brown spot, and soybean rust. Fungicides included: quinone outside inhibitors = azoxystrobin, fluoxastrobin, picoxystrobin, pyraclostrobin, trifloxystrobin; demethylation inhibitors = cyproconazole, difenoconazole, flutriafol, propiconazole, prothioconazole, tebuconazole, tetraconazole; methyl benzimidazole carbamates = thiophanate-methyl; multi-site mode of action = chlorothalonil; and succinate dehydrogenase inhibitors = fluxapyroxad.

**Table 1.** Coefficient of determination (R^2^) and *P*-values from linear and second order polynomial (quadratic) regression analyses for the relationship between foliar fungicide use and soybean yield losses due to foliar diseases from soybean growing states in the United States (α = 0.05) at national and regional (north and south) scales. A = relationship between annual total fungicide use (MT) and annual total production loss (1,000 MT) during 2005 to 2015. B = relationship between annual total fungicide use (g/ha) and annual yield loss (kg/ha) during 2005-2015. Northern states considered for this study included IL, IN, IA, KS, MI, MN, NE, ND, OH, PA, SD, and WI while southern states included AL, AR, DE, FL, GA, KY, LA, MD, MS, MO, NC, OK, SC, TN, TX, and VA.

| Geographic scale | A |  |  |  | B |  |  |  |
| --- | --- | --- | --- | --- | --- | --- | --- | --- |
|  | Linear |  | Quadratic |  | Linear |  | Quadratic |  |
| | $R^2$ | $P$-value | $R^2$ | $P$-value | $R^2$ | $P$-value | $R^2$ | $P$-value |
| National | 0.0683 | < 0.0001 | 0.1227 | < 0.0001 | 0.0012 | 0.5368 | 0.0016 | 0.7682 |
| North | 0.0924 | 0.0004 | 0.0975 | 0.0013 | 0.0864 | 0.0006 | 0.0888 | 0.0025 |
| South | 0.1671 | < 0.0001 | 0.2157 | < 0.0001 | 0.0279 | 0.0300 | 0.0305 | 0.0686 |

At a regional level (northern and southern), significant quadratic relationships were observed between total yield losses due to foliar diseases (1,000 MT) and total foliar fungicide use (MT) (Fig. 4C). For both regions, the total yield losses initially increased, reaching a plateau with increasing fungicide use. A stronger relationship was observed for the southern states compared to the northern states. When the losses (kg) and fungicide use (g) were considered on a per hectare basis, a significant quadratic relationship was observed for the northern states while no significant quadratic relationship was evident for the southern states (Fig. 4D). However, the linear relationship was significant and positive for southern states (Table 1).

### Relationship between annual soybean yield losses and annual fungicide use at the state level

Table 2 presents the coefficient of determination (R^2^) and corresponding *P*-values from linear and quadratic regression analyses for foliar fungicide use and soybean yield losses due to foliar diseases. A positive linear relationship was observed between total yield losses (1,000 MT) and total fungicide use (MT) for Florida (*y = 0.52x + 0.3*), Indiana (*y = 4.36x – 166.7*), Kentucky (*y = 0.65x – 7.2*), and Pennsylvania (*y = 6.49x + 21.3*) while a significant quadratic relationship was observed for Louisiana (*y = 0.001x^2^ + 0.16x +27.1*), Missouri (*y = 0.069x^2^ – 5.65x + 126.0*), and Virginia (*y = –0.030x^2^ + 1.64x – 4.1*). When losses (kg) and fungicide use (g) were considered on a per hectare basis, a negative linear relationship was observed for South Carolina (*y = –0.07x + 37.3*) while a significant quadratic relationship was evident for Indiana (*y = –0.613x^2^ + 38.9x – 551.3*), Louisiana (*y = 3E-06x^2^ + 0.35x + 61.3*) and Missouri (*y = 0.1473x^2^ – 6.07x + 67.7*).

**Table 2.** Coefficient of determination (R^2^) and corresponding *P*-values from linear and second order polynomial (quadratic) regression analyses for the relationship between foliar fungicide use and soybean yield losses due to foliar diseases from soybean growing states in the United States (α = 0.05). A = relationship between annual total fungicide use (MT) and annual total production loss (1,000 MT) during 2005 to 2015. B = relationship between annual total fungicide use (g/ha) and annual yield loss (kg/ha) during 2005 to 2015. Bold values indicate significant *P*-values and associated R^2^ values.

| Region | State | A |  |  |  | B |  |  |  |
| --- | --- | --- | --- | --- | --- | --- | --- | --- | --- |
|  |  | Linear |  | Quadratic |  | Linear |  | Quadratic |  |
| | | $R^2$ | $P$ | $R^2$ | $P$ | $R^2$ | $P$ | $R^2$ | $P$ |
| North | Illinois | 0.051 | 0.5037 | 0.051 | 0.8098 | 0.081 | 0.3967 | 0.101 | 0.6512 |
|  | Indiana | **0.484** | **0.0174** | 0.514 | 0.0558 | **0.421** | **0.0307** | **0.630** | **0.0188** |
|  | Iowa | 0.007 | 0.8127 | 0.007 | 0.9739 | 0.000 | 0.9593 | 0.001 | 0.9940 |
|  | Kansas | 0.004 | 0.8612 | 0.072 | 0.7426 | 0.000 | 0.9912 | 0.042 | 0.8420 |
|  | Michigan | 0.048 | 0.5163 | 0.073 | 0.7399 | 0.037 | 0.5726 | 0.054 | 0.8019 |
|  | Minnesota | 0.090 | 0.3698 | 0.328 | 0.2043 | 0.049 | 0.5091 | 0.232 | 0.3472 |
|  | Nebraska | 0.061 | 0.4626 | 0.162 | 0.4911 | 0.047 | 0.5196 | 0.165 | 0.4860 |
|  | North Dakota | 0.088 | 0.3763 | 0.192 | 0.4269 | 0.024 | 0.6467 | 0.081 | 0.7116 |
|  | Ohio | 0.107 | 0.3251 | 0.107 | 0.6346 | 0.086 | 0.3805 | 0.088 | 0.6903 |
|  | Pennsylvania | **0.442** | **0.0255** | 0.451 | 0.0911 | 0.310 | 0.0748 | 0.311 | 0.2256 |
|  | South Dakota | 0.008 | 0.7869 | 0.095 | 0.6698 | 0.029 | 0.6163 | 0.101 | 0.6543 |
|  | Wisconsin | 0.147 | 0.2435 | 0.283 | 0.2636 | 0.098 | 0.3482 | 0.397 | 0.1321 |
| South | Alabama | 0.318 | 0.0708 | 0.323 | 0.2102 | 0.011 | 0.7603 | 0.018 | 0.9289 |
|  | Arkansas | 0.165 | 0.2145 | 0.304 | 0.2352 | 0.163 | 0.2180 | 0.287 | 0.2583 |
|  | Delaware | 0.010 | 0.7678 | 0.049 | 0.8188 | 0.007 | 0.7943 | 0.068 | 0.7535 |
|  | Florida | **0.428** | **0.0287** | 0.436 | 0.1006 | 0.001 | 0.9169 | 0.024 | 0.9058 |
|  | Georgia | 0.307 | 0.0768 | 0.443 | 0.0960 | 0.077 | 0.4063 | 0.098 | 0.6606 |
|  | Kentucky | **0.428** | **0.0288** | 0.429 | 0.1062 | 0.267 | 0.1037 | 0.387 | 0.1409 |
|  | Louisiana | **0.859** | **<0.0001** | **0.868** | **0.0003** | **0.717** | **0.0010** | **0.717** | **0.0064** |
|  | Maryland | 0.009 | 0.7729 | 0.175 | 0.4627 | 0.004 | 0.8465 | 0.164 | 0.4888 |
|  | Mississippi | 0.106 | 0.3291 | 0.295 | 0.2466 | 0.038 | 0.5641 | 0.080 | 0.7162 |
|  | Missouri | **0.425** | **0.0297** | **0.592** | **0.0277** | **0.593** | **0.0055** | **0.847** | **0.0006** |
|  | North Carolina | 0.108 | 0.3244 | 0.112 | 0.6215 | 0.099 | 0.3463 | 0.122 | 0.5938 |
|  | Oklahoma | 0.068 | 0.4378 | 0.154 | 0.5130 | 0.021 | 0.6694 | 0.033 | 0.8743 |
|  | South Carolina | 0.064 | 0.4534 | 0.065 | 0.7617 | **0.364** | **0.0495** | 0.481 | 0.0724 |
|  | Tennessee | 0.003 | 0.8799 | 0.115 | 0.6135 | 0.032 | 0.5974 | 0.088 | 0.6898 |
|  | Texas | 0.176 | 0.1984 | 0.177 | 0.4571 | 0.002 | 0.8998 | 0.066 | 0.7599 |
|  | Virginia | 0.005 | 0.8361 | **0.530** | **0.0490** | 0.043 | 0.5401 | 0.045 | 0.8304 |

### Relationship between soybean yield losses and fungicide use over time

Table 3 presents the coefficient of variation (R^2^) and corresponding *P*-values from linear and quadratic regression analyses for foliar fungicide use and soybean yield losses due to foliar diseases for soybean growing states in the United States from 2005 to 2015. A positive linear relationship was observed between total yield losses due to foliar diseases (1,000 MT) and total foliar fungicide use (MT) in 2006 (*y = 2.87x + 16.7*) and 2010 (*y = 1.98x + 27.2*) while a significant quadratic relationship was observed in 2008 (*y = −0.003x^2^ + 2.39x – 4.6*) and 2015 (*y = −0.01x^2^ + 3.29x – 3.6*). When losses (kg) and fungicide use (g) were considered on a per hectare basis, neither linear nor quadratic relationships was significant for any given year between 2005 and 2015.

**Table 3.** Coefficient of determination (R^2^) and corresponding *P*-values from linear and second order polynomial (quadratic) regression analyses for the relationship between foliar fungicide use and soybean production/yield losses due to foliar diseases from soybean growing states in the United States during 2005 to 2015 period (α = 0.05). A = relationship between total fungicide use (MT) and total soybean production loss (1,000 MT) per state basis. B = relationship between total fungicide use (g/ha) and total yield loss (kg/ha) per state basis. Bold values indicate significant *P*-values and associated R^2^ values.

| Year | A |  |  |  | B |  |  |  |
| --- | --- | --- | --- | --- | --- | --- | --- | --- |
|  | Linear |  | Quadratic |  | Linear |  | Quadratic |  |
| | $R^2$ | $P$ | $R^2$ | $P$ | $R^2$ | $P$ | $R^2$ | $P$ |
| 2005 | 0.044 | 0.2814 | 0.093 | 0.2932 | 0.052 | 0.2431 | 0.091 | 0.3055 |
| 2006 | **0.141** | **0.0488** | 0.182 | 0.0814 | 0.000 | 0.9835 | 0.005 | 0.9331 |
| 2007 | 0.000 | 0.9574 | 0.039 | 0.6060 | 0.000 | 0.8857 | 0.003 | 0.9675 |
| 2008 | **0.234** | **0.0091** | **0.235** | **0.0354** | 0.004 | 0.7471 | 0.022 | 0.7608 |
| 2009 | 0.073 | 0.1633 | 0.207 | 0.0547 | 0.011 | 0.5978 | 0.027 | 0.7083 |
| 2010 | **0.176** | **0.0265** | 0.176 | 0.0892 | 0.039 | 0.3079 | 0.046 | 0.5584 |
| 2011 | 0.014 | 0.5439 | 0.045 | 0.5652 | 0.091 | 0.1196 | 0.147 | 0.1361 |
| 2012 | 0.085 | 0.1328 | 0.090 | 0.3086 | 0.048 | 0.2606 | 0.048 | 0.5376 |
| 2013 | 0.022 | 0.4533 | 0.053 | 0.5059 | 0.017 | 0.5022 | 0.044 | 0.5686 |
| 2014 | 0.043 | 0.2869 | 0.203 | 0.0588 | 0.008 | 0.6566 | 0.046 | 0.5554 |
| 2015 | 0.114 | 0.0795 | **0.302** | **0.0112** | 0.004 | 0.7464 | 0.074 | 0.3802 |

### Relationship between annual soybean production/yield and annual fungicide use at national and regional levels

Table 4 presents the coefficient of determination (R^2^) and corresponding *P*-values for the linear and quadratic relationships between soybean production/yield and fungicide use at national and regional level. Analysis of the national level dataset revealed a significant quadratic relationship between total annual soybean production (1,000 MT) and total annual foliar fungicide use (MT) during the period between 2005 and 2015 (Fig. 5A). The total soybean production initially increased, reached a plateau, and subsequently decreased with increasing fungicide use. Although a significant quadratic relationship was observed between soybean yield (kg) and fungicide use (g) on a per hectare basis, the yield initially decrease, reaches a plateau, and eventually increase slightly with increasing fungicide use (Fig. 5B)

**Fig 5.**
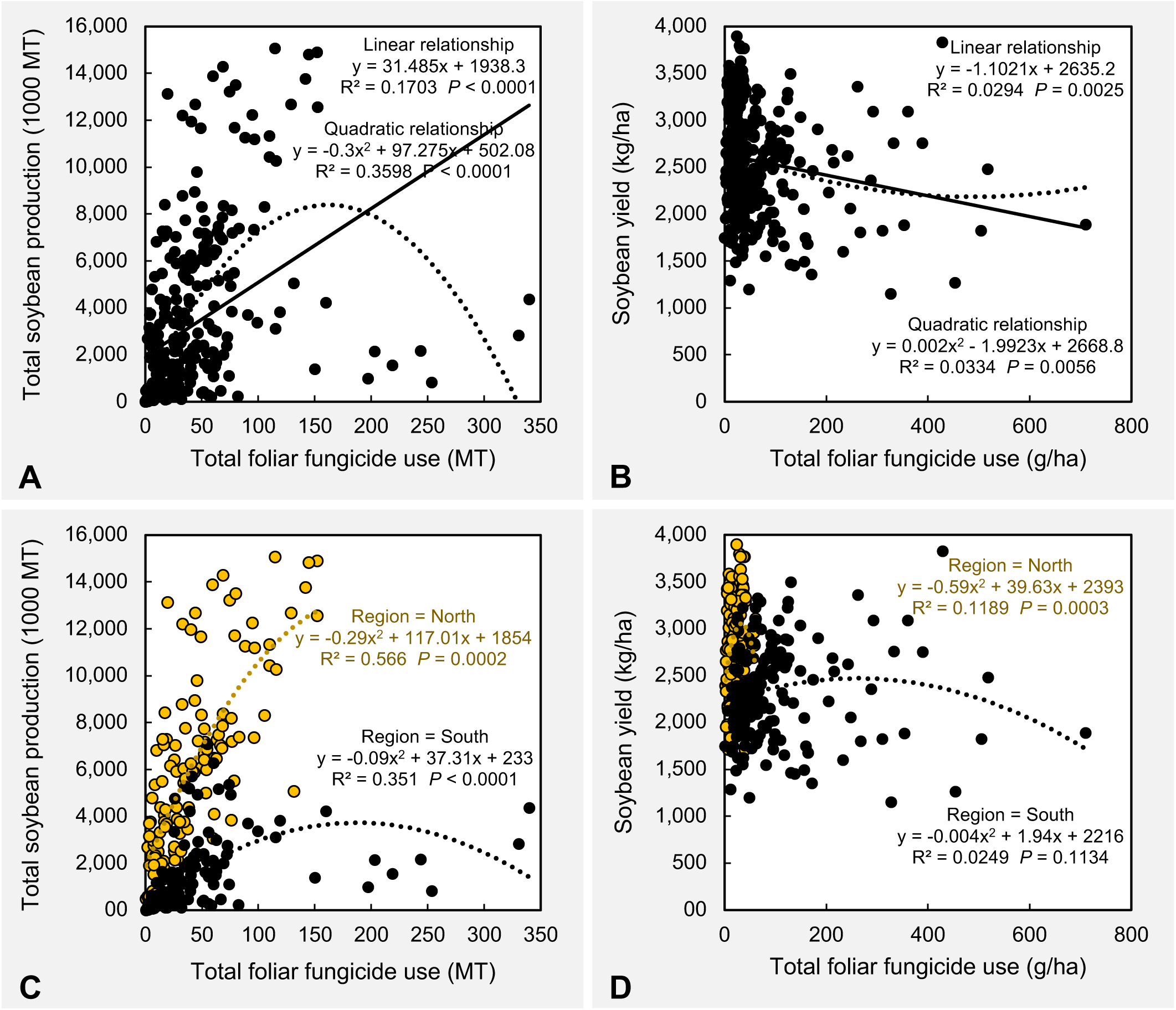
Scatter plots presenting the national scale linear/quadratic relationship between (A) total soybean production (1,000 MT) and total foliar fungicide use (MT), (B) soybean yield (kg/ha) and per hectare total foliar fungicide use (g), and regional scale quadratic relationship between (C) total soybean production (1,000 MT) and foliar fungicide use (MT), and (D) soybean yield (kg/ha) and per hectare total foliar fungicide use (g). Each data point represents a state in a given year. All plots contain data during 2005 to 2015 from 28 soybean growing states including AL, AR, DE, FL, GA, IA, IL, IN, KS, KY, LA, MD, MI, MN, MO, MS, NC, ND, NE, OH, OK, PA, SC, SD, TN, TX, VA, and WI. Fungicides included: quinone outside inhibitors = azoxystrobin, fluoxastrobin, picoxystrobin, pyraclostrobin, trifloxystrobin; demethylation inhibitors = cyproconazole, difenoconazole, flutriafol, propiconazole, prothioconazole, tebuconazole, tetraconazole; methyl benzimidazole carbamates = thiophanate-methyl; multi-site mode of action = chlorothalonil; and succinate dehydrogenase inhibitors = fluxapyroxad.

**Table 4.** Coefficient of determination (R^2^) and *P*-values from linear and second order polynomial (quadratic) regression analyses for the relationship between foliar fungicide use and soybean production/yield from soybean growing states in the United States (α = 0.05) at national and regional (north and south) scales. A = relationship between annual total fungicide use (MT) and annual total soybean production (1,000 MT) during 2005 to 2015. B = relationship between annual total fungicide use (g/ha) and annual yield (kg/ha) during 2005-2015. Northern states considered for this study included IL, IN, IA, KS, MI, MN, NE, ND, OH, PA, SD, and WI while southern states included AL, AR, DE, FL, GA, KY, LA, MD, MS, MO, NC, OK, SC, TN, TX, and VA.

| Geographic scale | A |  |  |  | B |  |  |  |
| --- | --- | --- | --- | --- | --- | --- | --- | --- |
|  | Linear |  | Quadratic |  | Linear |  | Quadratic |  |
| | $R^2$ | $P$-value | $R^2$ | $P$-value | $R^2$ | $P$-value | $R^2$ | $P$-value |
| National | 0.1703 | < 0.0001 | 0.3598 | < 0.0001 | 0.0294 | 0.0025 | 0.0334 | 0.0056 |
| North | 0.5509 | < 0.0001 | 0.5660 | 0.0002 | 0.0558 | 0.0064 | 0.1189 | 0.0003 |
| South | 0.2016 | < 0.0001 | 0.3506 | < 0.0001 | 0.0010 | 0.6724 | 0.0249 | 0.1134 |

At a regional level, a significant quadratic relationship was observed between total annual soybean production (1,000 MT) and total annual foliar fungicide use (MT) for both the northern and southern states (Fig. 5C). However, the magnitude of the relationship was stronger in the northern compared to the southern states. When the annual soybean yield (kg) and annual fungicide use (g) were considered on a per hectare basis, a significant quadratic relationship was observed for northern states while no significant linear or quadratic relationship was evident for the southern states (Fig. 5D). The quadratic curve fitted for the north showed that soybean yield increase at the beginning, reached a plateau (at approximately 35 g/ha), and subsequently decreased with increasing fungicide use.

### Relationship between annual soybean production/yield and annual fungicide use at state level

Table 5 presents coefficient of determination (R^2^) and corresponding *P*-values from linear and quadratic regression analyses for soybean production/yield and foliar fungicide use for soybean growing states in the U.S. A positive linear relationship was observed between total soybean production (1,000 MT) and total foliar fungicide use (MT) for Ohio (*y = 24.8x + 4,796.6*) while significant quadratic relationship was evident for Alabama (*y = −0.387x^2^ + 24.9x + 77.4*), Arkansas (*y = −0.019x^2^ + 12.1x + 2,547*), Florida (*y = −14.689x^2^ + 41.2x + 4.9*), Georgia (*y = - 0.149x^2^ + 14.6x + 52.8*), Illinois (*y = 0.632x^2^ – 100.6x + 14,835*), Kentucky (*y = –0.986x^2^ + 88.2x + 108.5*), Louisiana (*y = 0.016x^2^ – 0.12x + 919.7*), Mississippi (*y = −0.057x^2^ + 24.5x + 960.5*), North Dakota (*y = –0.1x^2^ + 30.2x + 2,899.5*), Pennsylvania (*y = –2.777x^2^ + 58.5x + 399.9*), and Tennessee (*y = –0.149x^2^ + 40.1x + 154.5*). When soybean yield (kg) and fungicide use (g) were considered on a per hectare basis, a positive linear relationship was observed for Pennsylvania (*y = 8.678x + 2,718.7*), while a significant quadratic relationship was observed for Alabama (*y = – 0.096x^2^ + 26.5x + 978.4*), Arkansas (*y = –0.029x^2^ + 14.7x + 1,532.8*), Kentucky (*y = –0.414x^2^ + 55.8x + 1,170*), and Mississippi (*y = –0.039x^2^ + 18.7x + 1,468.4*).

**Table 5.** Coefficient of determination (R^2^) and *P* values from linear and second order polynomial (quadratic) regression analyses for the relationship between foliar fungicide use and soybean production/yield from soybean growing states in the United States (α = 0.05). A = relationship between annual total fungicide use (MT) and annual total soybean production (1,000 MT) during 2005 to 2015. B = relationship between annual total fungicide use (g/ha) and annual yield (kg/ha) during 2005 to 2015. Bold values indicate significant *P*-values and associated R^2^ values.

| Region | State | A |  |  |  | B |  |  |  |
| --- | --- | --- | --- | --- | --- | --- | --- | --- | --- |
|  |  | Linear |  | Quadratic |  | Linear |  | Quadratic |  |
| | | $R^2$ | $P$ | $R^2$ | $P$ | $R^2$ | $P$ | $R^2$ | $P$ |
| North | Illinois | 0.186 | 0.1853 | **0.574** | **0.0329** | 0.229 | 0.1363 | 0.413 | 0.1184 |
|  | Indiana | 0.350 | 0.0550 | 0.415 | 0.1171 | 0.342 | 0.0588 | 0.344 | 0.1848 |
|  | Iowa | 0.016 | 0.7144 | 0.026 | 0.8994 | 0.000 | 0.9414 | 0.183 | 0.4465 |
|  | Kansas | 0.168 | 0.2103 | 0.296 | 0.2462 | 0.051 | 0.5042 | 0.127 | 0.5819 |
|  | Michigan | 0.105 | 0.3307 | 0.289 | 0.2563 | 0.278 | 0.0953 | 0.393 | 0.1360 |
|  | Minnesota | 0.084 | 0.3883 | 0.275 | 0.2756 | 0.111 | 0.3161 | 0.333 | 0.1974 |
|  | Nebraska | 0.283 | 0.0922 | 0.296 | 0.2459 | 0.186 | 0.1853 | 0.223 | 0.3643 |
|  | North Dakota | **0.578** | **0.0066** | **0.602** | **0.0249** | 0.131 | 0.2732 | 0.153 | 0.5143 |
|  | Ohio | **0.364** | **0.0495** | 0.501 | 0.0619 | 0.270 | 0.1013 | 0.387 | 0.1411 |
|  | Pennsylvania | **0.539** | **0.0101** | **0.632** | **0.0183** | **0.430** | **0.0285** | 0.479 | 0.0735 |
|  | South Dakota | 0.075 | 0.4139 | 0.249 | 0.3175 | 0.002 | 0.8951 | 0.101 | 0.6526 |
|  | Wisconsin | 0.027 | 0.6261 | 0.110 | 0.6268 | 0.023 | 0.6513 | 0.049 | 0.8197 |
| South | Alabama | **0.458** | **0.0222** | **0.539** | **0.0451** | 0.038 | 0.5635 | **0.534** | **0.0472** |
|  | Arkansas | **0.636** | **0.0033** | **0.719** | **0.0062** | 0.672 | 0.0020 | **0.737** | **0.0048** |
|  | Delaware | 0.200 | 0.1675 | 0.353 | 0.1756 | 0.104 | 0.3324 | 0.330 | 0.2018 |
|  | Florida | **0.472** | **0.0194** | **0.712** | **0.0069** | 0.355 | 0.0531 | 0.415 | 0.1170 |
|  | Georgia | 0.163 | 0.2178 | **0.602** | **0.0250** | 0.006 | 0.8201 | 0.116 | 0.6111 |
|  | Kentucky | **0.519** | **0.0124** | **0.633** | **0.0181** | **0.475** | **0.0274** | **0.619** | **0.0343** |
|  | Louisiana | **0.587** | **0.0060** | **0.604** | **0.0247** | 0.256 | 0.1120 | 0.429 | 0.1062 |
|  | Maryland | 0.101 | 0.3397 | 0.504 | 0.0605 | 0.086 | 0.3832 | 0.405 | 0.1251 |
|  | Mississippi | **0.710** | **0.0022** | **0.712** | **0.0128** | **0.576** | **0.0109** | **0.582** | **0.0471** |
|  | Missouri | 0.062 | 0.4585 | 0.285 | 0.2617 | 0.074 | 0.4192 | 0.095 | 0.6694 |
|  | North Carolina | 0.135 | 0.2968 | 0.254 | 0.3593 | 0.038 | 0.5866 | 0.041 | 0.8642 |
|  | Oklahoma | 0.223 | 0.1428 | 0.231 | 0.3488 | 0.026 | 0.6324 | 0.296 | 0.2449 |
|  | South Carolina | 0.052 | 0.5001 | 0.140 | 0.5466 | 0.193 | 0.1769 | 0.206 | 0.3971 |
|  | Tennessee | **0.603** | **0.0083** | **0.722** | **0.0113** | 0.189 | 0.1814 | 0.406 | 0.1245 |
|  | Texas | 0.006 | 0.8140 | 0.368 | 0.1600 | 0.064 | 0.4534 | 0.350 | 0.1785 |
|  | Virginia | 0.021 | 0.6684 | 0.462 | 0.0836 | 0.081 | 0.3938 | 0.154 | 0.5133 |

### Relationship between soybean production/yield and fungicide use over time

Table 6 presents coefficient of variation (R^2^) and corresponding *P*-values from linear and quadratic regression analyses for foliar fungicide use and soybean production/yield for soybean growing states in the United States from 2005 to 2015. Except for 2007, a significant quadratic relationship was observed between total soybean production (1,000 MT) and total foliar fungicide use (MT) for all other years (2005: *y = 2.271x^2^ + 27.2x + 906.5*; 2006: *y = 0.381x^2^ + 127.3x + 672*; 2008: *y = 0.199x^2^ + 85.4x + 251.1*; 2009: *y = – 0.597x^2^ + 125.6x + 255.1*; 2010: *y = 0.594x^2^ + 63.9x + 828*; 2011: *y = 0.489x^2^ + 20.8x + 800*; 2012: *y = – 0.731x^2^ + 139.1x – 906.5*; 2013: *y = – 0.129x^2^ + 97.3x – 635*; 2014: *y = – 0.298x^2^ + 109.1x – 255.2*; 2015: *y = – 0.301x^2^ + 98x + 126.6*). When soybean yield (kg) and foliar fungicide use (g) were considered on a per hectare basis, a significant quadratic relationship was only observed for 2005 (*y = 0.037x^2^ – 13.8x + 2,826.2*), 2007 (*y = 0.006x^2^ – 4.9x + 2,608*), 2010 (*y = 0.038x^2^ – 15.2x + 2,991.5*), and 2011 (*y = – 0.059x^2^ + 1.6x + 2,583.4*).

**Table 6.** Coefficient of determination (R^2^) and corresponding *P*-values from linear and second order polynomial (quadratic) regression analyses for the relationship between foliar fungicide use and soybean production/yield from soybean growing states in the United States during 2005 to 2015 period (α = 0.05). A = relationship between total fungicide use (MT) and total soybean production (1,000 MT) per state basis. B = relationship between total fungicide use (g/ha) and yield (kg/ha) per state basis. Bold values indicate significant *P*-values and associated R^2^ values.

| Year | A |  |  |  | B |  |  |  |
| --- | --- | --- | --- | --- | --- | --- | --- | --- |
|  | Linear |  | Quadratic |  | Linear |  | Quadratic |  |
|  | R<sup>2</sup> | P | R<sup>2</sup> | P | R<sup>2</sup> | P | R<sup>2</sup> | P |
| 2005 | **0.518** | **<0.0001** | **0.564** | **<0.0001** | **0.276** | **0.0041** | **0.371** | **0.0030** |
| 2006 | **0.407** | **0.0003** | **0.408** | **0.0014** | 0.093 | 0.1155 | 0.128 | 0.1798 |
| 2007 | 0.004 | 0.7410 | 0.028 | 0.6966 | **0.275** | **0.0042** | **0.375** | **0.0028** |
| 2008 | **0.474** | **<0.0001** | **0.476** | **0.0003** | 0.108 | 0.0879 | 0.127 | 0.1819 |
| 2009 | **0.355** | **0.0010** | **0.512** | **0.0002** | 0.048 | 0.2604 | 0.103 | 0.2570 |
| 2010 | **0.404** | **0.0003** | **0.409** | **0.0014** | **0.399** | **0.0003** | **0.407** | **0.0015** |
| 2011 | **0.634** | **<0.0001** | **0.669** | **<0.0001** | **0.233** | **0.0093** | **0.258** | **0.0239** |
| 2012 | **0.578** | **<0.0001** | **0.585** | **<0.0001** | 0.021 | 0.4670 | 0.046 | 0.5554 |
| 2013 | **0.740** | **<0.0001** | **0.745** | **<0.0001** | 0.035 | 0.3370 | 0.049 | 0.5317 |
| 2014 | **0.196** | **0.0182** | **0.583** | **<0.0001** | 0.077 | 0.1518 | 0.128 | 0.1797 |
| 2015 | 0.103 | 0.0954 | **0.440** | **0.0007** | 0.018 | 0.4948 | 0.185 | 0.0780 |

### Factor analysis with mixed data (FAMD)

When FAMD was performed for foliar fungicide use, the variance maximizing data point (a data point = total fungicide use in a particular year for a given state) distribution in the factor map did not show a clear clustering pattern based upon state, year, and yield zone. However, a clear clustering was observed based upon region, harvest zone, and production zone (Fig. 6). Factor maps for both harvest and production zones showed that harvest/production zone 1 distantly clusters from harvest/production zone 4 while harvest/production zones 1 and 2 clustered in close proximity in the factor map.

**Fig 6.**
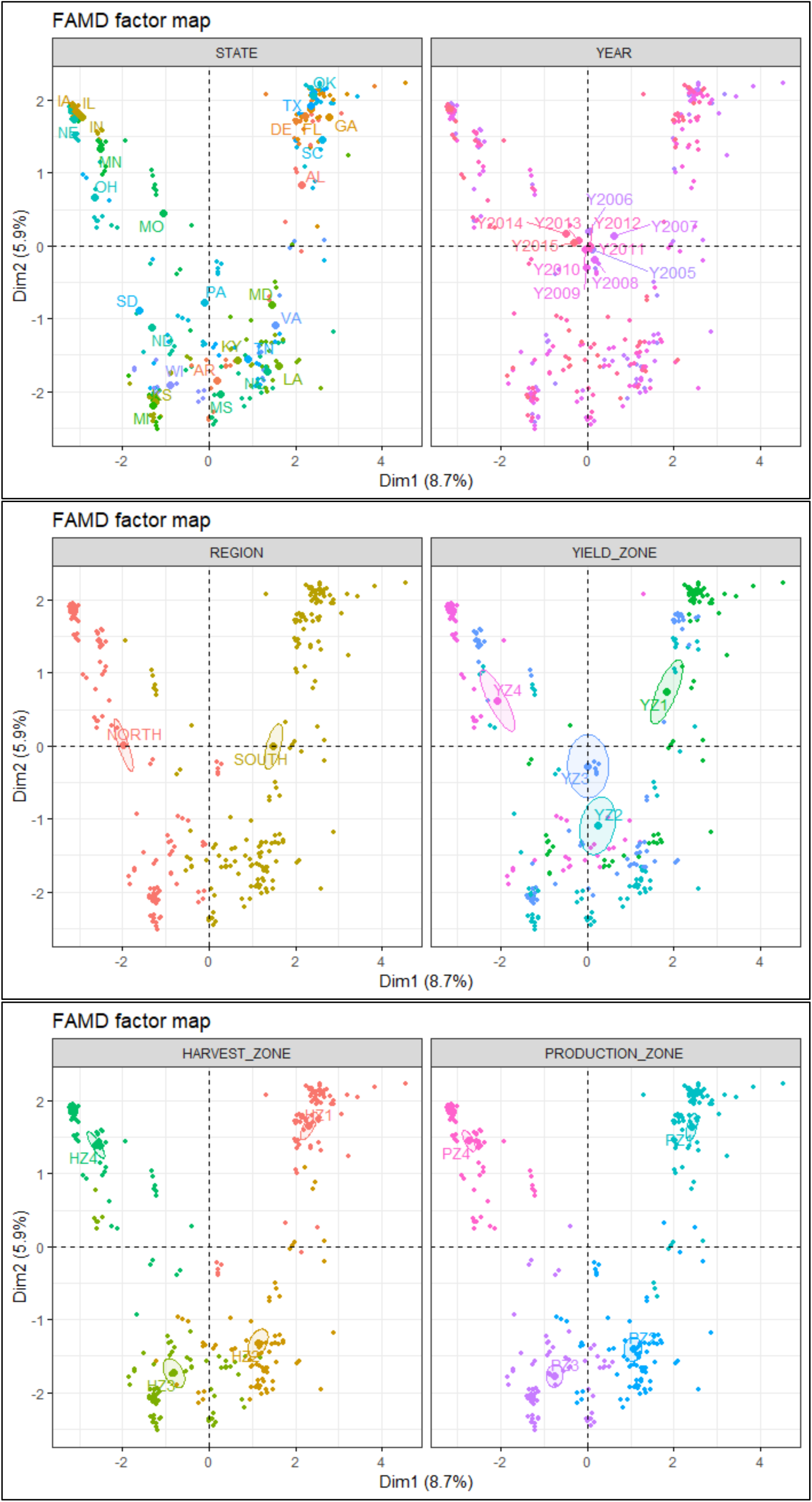
FAMD factor maps obtained from the factor analysis with mixed data approach (FAMD analysis), showing the variance maximizing distribution pattern of data points (n = 308, each data point represent foliar fungicide use in g/ha) in the map space with their clustering patterns based upon state (n = 28), year (n = 11), region (n = 2), and yield/harvest/production zones (n = 4 in each case). Yield/Harvest/Production zones = represent four levels (zone 1 to 4) based on the quartiles within a database containing 308 yield (kg/ha)/harvest area (ha)/production (MT) data points (308 = 11 years × 28 states). Within this database, data points from the minimum to the first quartile were classified as zone 1. Similarly, data points from the first quartile to median, median to the third quartile, and > third quartile were respectively classified as zones 2, 3, and 4. Foliar fungicides included: quinone outside inhibitors = azoxystrobin, fluoxastrobin, picoxystrobin, pyraclostrobin, trifloxystrobin; demethylation inhibitors = cyproconazole, difenoconazole, flutriafol, propiconazole, prothioconazole, tebuconazole, tetraconazole; methyl benzimidazole carbamates = thiophanate-methyl; multi-site mode of action = chlorothalonil; and succinate dehydrogenase inhibitors = fluxapyroxad (effective against anthracnose, Cercospora leaf blight (purple seed stain), frogeye leaf spot, Rhizoctonia aerial blight, Sclerotinia stem rot (White mold), Septoria brown spot, and soybean rust)

### Analysis of variance (ANOVA)

ANOVA showed significant main effect of yield zone (*P* = 0.0003), harvest zone (*P* < 0.0001), and production zone (*P* < 0.0001) on foliar fungicide use. With respect to yield zone, the foliar fungicide use (g/ha) in yield zone 1 was significantly greater than that of yield zones 2, 3, and 4 (Fig. 7). In case of harvest zone, the foliar fungicide use in zones 1 and 2 were significantly greater compared to those of zones 3 and 4. Foliar fungicide use in harvest zone 3 was significantly greater than that of zone 4. Trends observed for production zone were same as with harvest zone.

**Fig 7.**
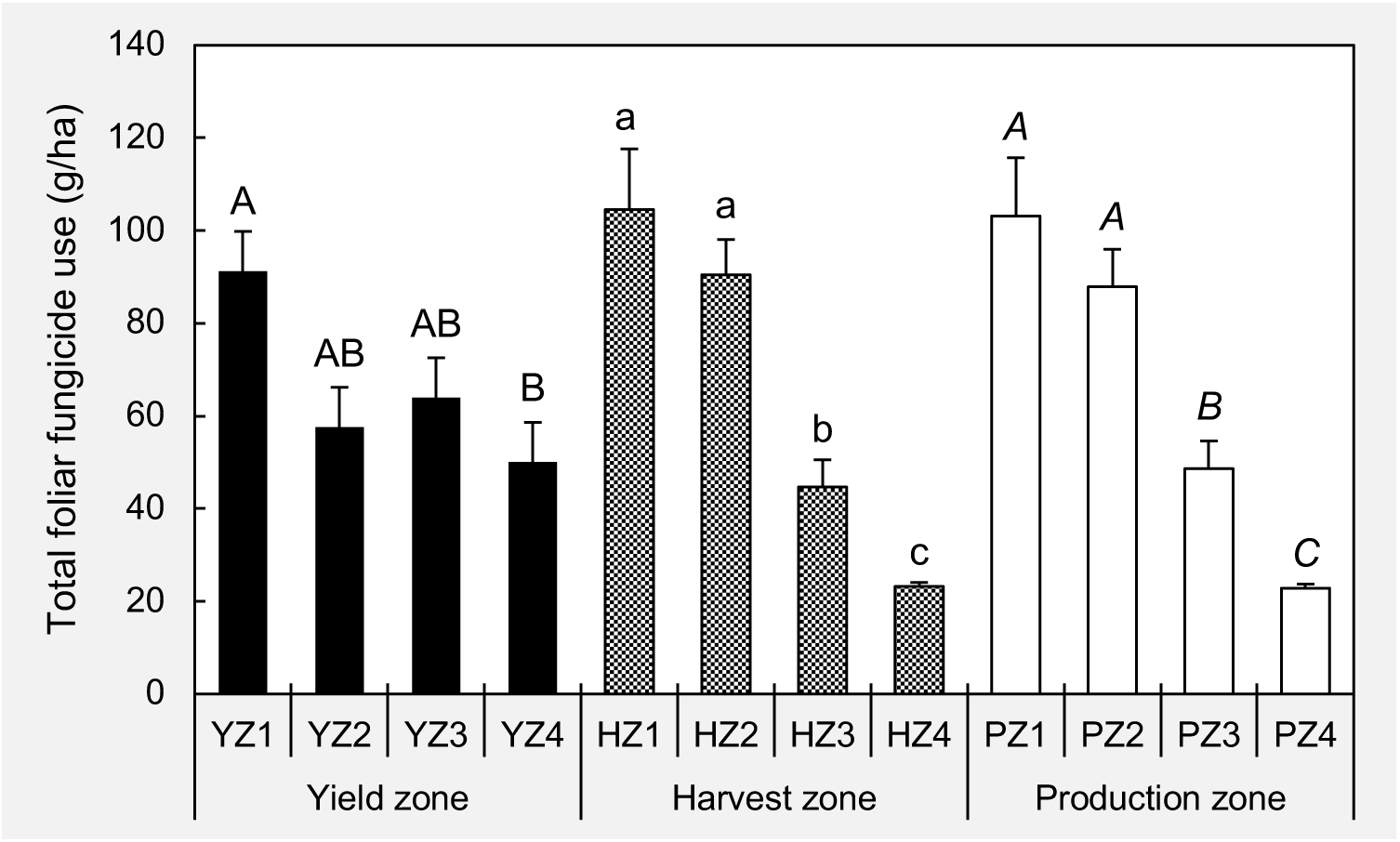
Comparison of the mean per hectare foliar fungicide use (in g) among yield/harvest/production zones. Treatment means with different letter designations (within each zone type) are significantly different at α = 0.05 based on the adjustment for multiple comparison using Tukey-Kramer test. Error bars represent standard errors. Foliar fungicides included: quinone outside inhibitors = azoxystrobin, fluoxastrobin, picoxystrobin, pyraclostrobin, trifloxystrobin; demethylation inhibitors = cyproconazole, difenoconazole, flutriafol, propiconazole, prothioconazole, tebuconazole, tetraconazole; methyl benzimidazole carbamates = thiophanate-methyl; multi-site mode of action = chlorothalonil; and succinate dehydrogenase inhibitors = fluxapyroxad (effective against anthracnose, Cercospora leaf blight (purple seed stain), frogeye leaf spot, Rhizoctonia aerial blight, Sclerotinia stem rot (White mold), Septoria brown spot, and soybean rust)

## DISCUSSION

The current paper seeks to understand the patterns of foliar fungicide use and its relationship with soybean yield losses due to fungal pathogens (targets of fungicides considered in the study) at broader geographic (national/regional/state) and temporal scales. The trends that we see at such scales may or may not necessarily reflect/represent what each individual soybean farmer would have experienced at a farm scale. For example, the lack of a strongly negative relationship between yield losses and fungicide use at the state level does not necessarily mean that all soybean farmers within that particular state did not experience a negative relationship between fungicide use and soybean yield losses at their individual farm level. In other words, with the results we present here, we cannot simply extrapolate the individual farmer experience in relation to his fungicide use and yield losses profiles. As such, this paper does not necessarily seek to facilitate the fungicide application decision making at the individual farm level, rather trying to provide some insights on fungicide use patterns and their degree of utility in terms of reducing foliar disease associated yield losses at a broader geographic scale.

Use of foliar fungicides has been a major strategy to manage fungal pathogens in agricultural cropping systems following the green revolution. Fungicide usage has increased over the past decade especially in soybean production systems. Findings of the current study revealed that the foliar fungicide usage in the U.S. increased by 116% (on a per unit area basis: g/ha) and 260% (on a total usage basis: MT) from 2005 to 2015. Fungicide use was greatest in 2007, which was a year with more widespread soybean rust outbreaks on a national level and the first year that soybean rust moved into the upper Midwest through Texas to Iowa [36]. Furthermore, 2007 was the only year to date that Iowa reported observing the disease [36]. A similar situation occurred in 2009, where an increased incidence of soybean rust was reported. For example, Alabama, Georgia, Mississippi, and Tennessee reported the greatest number of counties with soybean rust [36]. Moreover, on a national basis, more counties/parishes were observed to contain soybean rust during 2009 than any other year [36]. Additionally, 2009 was an exceptionally wet year particularly in the southern U.S., leading to more foliar diseases [36]. All these factors could have specifically contributed to the greater foliar fungicide use in 2009.

The regional level data revealed that foliar fungicide use (total in MT as well and per hectare basis in g) was greater in the southern states compared to the northern states despite the greater land use for soybean production in the northern states. The greater per hectare fungicide use in the south may be due to several reasons. In general, this region has an extended period of soybean planting (March to June) and a prolonged period of disease conducive conditions (warmer and wetter for a longer period of time) compared to the northern U.S. Along with that, soybean rust was first detected in the contiguous U.S. in November 2004 [37] and fungicides were the main method of managing the disease. Even though soybean rust has not posed a major yield loss threat since the initial observation [38], fungicide applications in specific years have likely been driven by the presence of the disease. Lastly, based on observations by Extension specialists, a greater percentage (60-65 %) of southern U.S. acres likely receives at least one fungicide application at a specific growth stage as an automatic application in the absence of diseases.

The significant quadratic relationship observed at the national level between total yield losses (in 1,000 MT) due to foliar diseases and total foliar fungicide use (MT) indicated that losses initially increased at a decreasing rate, reached a plateau, and eventually decreased at an increasing rate with increasing use of fungicide. This was observed in both northern and southern states for the relationship between total yield losses due to foliar diseases and total foliar fungicide use. The positive relationship observed between yield losses and fungicide use in the initial phase of the curve (at both national and regional levels) could occur as a result of the poor application timing practices of fungicide with the intention of saving the crop under severe disease pressure, particularly after passing the economic injury level. The application timing greatly affects the effectiveness of a fungicide in terms of its ability to suppress the severity of a disease and associated yield losses [19, 26, 39, 40]. For instance, applying a suitable fungicide after the establishment of a disease for diseases that include frogeye leaf spot [39] and soybean rust [41] could still result in significant yield losses.

Prophylactic application of foliar fungicides can significantly increase production costs, and subsequently suppress profitability particularly when diseases are absent or are present at low levels [42]. In the current study, we observed that the vast majority of the states have used a greater amount of preventive fungicides as compared to curative fungicide at the temporal scale. If the application of a preventive fungicide was not made at the suggested growth stage based on plant phenology, such applications may not provide a scenario whereby a reduction in the potential yield losses associated with a given disease were met. Poor fungicide application practices may contribute to a positive relationship between fungicide use and yield losses.

Additionally, reduced fungicide efficiency due to a variety of factors such as unfavorable environmental conditions and automatic fungicide application on disease-resistant soybean cultivars can result in a positive relationship between fungicide use and yield losses. For example, compared to the control, application of benomyl at different application timings based on growth stage did not significantly reduce frogeye leaf spot severity or associated grain yield loss on resistant soybean genotypes, although significant disease severity and yield loss reductions were observed with susceptible soybean genotypes [39]. Resistance within the targeted pathogen population to the active ingredient contained in the applied fungicide/s could also contribute to a positive relationship between fungicide use and yield losses [28, 29, 31–33]. Furthermore, fungicides are applied with self-propelled, pull type, or aerial spray applicators in the U.S. Ground applicators create wheel-tracks in the soybean crop, which reduce yield particularly when made during the reproductive growth stages [43]. This also can contribute to positive correlation between fungicide use and soybean yield losses.

When yield losses and fungicide use were considered on a per hectare basis at both the national and regional levels, the relationships were non-significant, with the exception of the significant quadratic relationship between per hectare fungicide use and losses due to foliar diseases in the northern states. The non-significant relationships could partly be due to the application of fungicide in the absence of disease, under low disease pressure or when applied to a disease-resistant cultivar. In fact, several experimental studies regarding the prophylactic application of foliar fungicides in the absence of disease revealed non-significantly different yield between treated and non-treated situations [20–22]. These results suggest that the application of foliar fungicides may be more beneficial in the presence of disease or situations where there is a high probability of disease occurrence. For instance, experimental data revealed that farmers in the north central region do not often need fungicide applications to manage foliar diseases [23]. However, in many cases, increasing the number of inputs including a fungicide application has become part of a farmer’s soybean management system in their search for maximizing yield and subsequent profitability [42].

Analyses conducted at the state level showed no significant linear or quadratic relationship between soybean production/yield losses and foliar fungicide use for a vast majority of the states. Although a significant linear relationship between total yield losses (in 1,000 MT) due to foliar diseases and total foliar fungicide use (MT) was observed for Florida, Indiana, Kentucky, and Pennsylvania, the relationship was positive. Similarly, although the quadratic relationship between the two variables were significant for Louisiana and Missouri, the direction of the relationship was positive. Furthermore, when the losses and fungicide use were considered on a per hectare basis, a significant quadratic relationship was evident between them for Louisiana and Missouri. However, the direction of the relationship was positive. Therefore, at the state level, our findings do not provide strong statistical evidence to support the usefulness of foliar fungicide application to mitigate foliar disease-associated soybean yield losses. Nevertheless, a significant quadratic relationship was observed between total soybean production and total foliar fungicide use (MT) for Alabama, Arkansas, Florida, Georgia, Kentucky, Louisiana, Mississippi, North Dakota, Pennsylvania, and Tennessee. Quadratic curves for each of these states showed that increased fungicide use contributed to increased yield, although this reached a plateau where additional fungicide did not add additional yield. These relationships showed that there may be some benefit for fungicide use in soybean production systems.

Results from the factor analysis with mixed data (FAMD) showed clear distinction between yield/harvest/production zone 1 and 4 based on foliar fungicide use, suggesting contrasting fungicide use differences between these zones. The current observation was further confirmed by ANOVA results where mean use in zone 1 was significantly greater compared to zone 4. In general, results showed that the mean per hectare foliar fungicide use was significantly greater in low yield/harvest/production zones while the use was lower in high yield/harvest/production zones. However, it may be possible that soybean farmers in low yield/harvest/production zones tend to apply foliar fungicides based on a perceived yield benefit as the result of an application made at a specific growth stage, rather than based upon disease observations or soybean cultivar disease tolerance. In fact, previous studies suggested that yield increases can occur following foliar fungicide application irrespective of the presence/absence of diseases [7, 15, 23–25, 44–46]. The yield response in the absence of disease has been partly attributed to the physiological changes that have been reported to occur in the plants following fungicide application with certain chemistries [14]. Increased yield in response to some fungicides such as QoIs have been observed even in the absence of foliar diseases due to their non-fungicidal physiological changes in, for example, soybean [22, 47, 48], wheat, and barley [49–51]. Some of these plant physiological changes include increased leaf greenness, chlorophyll content, photosynthetic rates, and water use efficiency, as well as delayed senescence [44, 46, 49, 50, 52]. Previous studies also reported that foliar application of pyraclostrobin enhance the growth, nitrogen assimilation, and yield of soybean [53] and wheat [54, 55]. Therefore, as revealed by the current study, it appeared that the farmers in the historically low yield/harvest/production zones tend to use foliar fungicide applications with the expectation of a yield increase.

In the current study, it was not possible to determine the relationship between yield losses caused by a single disease and the amount of a labeled fungicide used to control that disease. This was because each fungicide considered in this study may effectively control more than one disease. For instance, QoI fungicides can be used to manage anthracnose (*Colletotrichum truncatum*), Cercospora leaf blight (*Cercospora kikuchii*), frogeye leaf spot; pod and stem blight (*Diaporthe phaseolorum*); Rhizoctonia aerial blight (*Rhizoctonia solani*), and Septoria brown spot [6, 7, 56, 57]. Based on the manner in which the information in the fungicide use database is provided, there is no way to tell what the fungicide specifically targeted. Therefore, relationships between total yield losses caused by all foliar diseases and total concentration of foliar fungicide used were considered for this study.

Although we have previously estimated soybean yield losses due to various diseases for the period between 1996 and 2015 [58], the corresponding annual state-level foliar fungicide use estimates were not available for the entire period in the Pesticide National Synthesis Project database (https://water.usgs.gov/nawqa/pnsp/usage/maps/county-level/StateLevel/HighEstimate_AgPestUsebyCropGroup92to16.txt). Therefore, the foliar fungicides used between 2005 and 2015 were considered for the current study. With the data used for this study, it was not possible to conduct a realistic economic analysis to determine whether fungicide application was cost effective. Unless there is an appropriate control for comparison, one could not determine the economic yield savings as a result of fungicides applied. Moreover, it is likely that the physical yield losses could have potentially been greater if fungicides were not applied. In addition, the fungicide database only contains information regarding the use of active ingredients and does not include such information as to whether or not a particular active ingredient was applied as a stand-alone fungicide product or in the form of a pre-mixture of more than one chemical. Based on the commercial product and company, the same active ingredient can be marketed under several different trade names and in some cases the products can be priced differently depending on retail outfit. Annual fluctuations as well as locational variations in fungicide application cost (i.e., aerial application versus ground application) and soybean commodity price also are contributing factors as to why a comprehensive economic analysis is less realistic.

In summary, we did not observe a strongly negative relationship between foliar fungicide use (g) and yield losses (Kg) per unit area (ha) at national, regional, and state levels. However, the observed positive impact of foliar fungicide use on total soybean production (1,000 MT) and yield (kg/ha) at national, regional, and state levels indicated the possible benefit of foliar fungicide application to produce greater soybean yield. Nevertheless, we cannot simply extrapolate the individual farmer experience in relation to his/her fungicide use and foliar disease associated yield losses (and total soybean production too) based on the trends that we observed in this paper at a broader spatial scale (national/regional/state). As such, the specific content of this paper may not be strictly useful to facilitate the fungicide application decision making at individual farm level. However, we suggest that farmers should not rely on fungicides as the sole management strategy to manage foliar diseases in soybean. Instead, location specific best management practices such as optimum maturity group, planting date, seeding rate, row spacing, crop rotation, fertilizer, field history as it relates to disease incidence, and irrigation regime as well as use of genetic resistance should be emphasized to decrease the probability of disease incidence. When necessary, farmers should make informed decisions as to the use of foliar fungicides with special emphasis on application timing (disease susceptible plant growth stage). In conclusion, rather than using fungicides as a routine practice, farmers should treat foliar fungicides as an integral component of a sound integrated pest management system.

## ACKNOWLEDGEMENTS

We thank the United Soybean Board for support of the soybean yield losses estimates. This project was also supported by the USDA National Institute of Food and Federal Appropriations under Project PEN04660 and Accession number 1016474. We also would like to express our thanks and gratitude to all Extension plant pathologist and soybean disease experts who contributed observations on soybean losses in their respective states over the years.

